# MAPK15 PROTECTS AGAINST THE DEVELOPMENT OF METABOLIC DYSFUNCTION-ASSOCIATED STEATOTIC LIVER DISEASE

**DOI:** 10.1101/2024.12.23.630083

**Authors:** Giovanni Inzalaco, Sara Gargiulo, Denise Bonente, Lisa Gherardini, Lorenzo Franci, Nicla Lorito, Serena Del Turco, Danilo Tatoni, Tiziana Tamborrino, Eugenio Bertelli, Romina D’Aurizio, Maria Grazia Andreassi, Giuseppina Basta, Amalia Gastaldelli, Andrea Morandi, Virginia Barone, Mario Chiariello

## Abstract

Accumulation of lipids in the liver characterizes metabolic dysfunction-associated steatotic liver disease (MASLD), the most prevalent chronic liver disease worldwide. As liver injury progresses to metabolic dysfunction-associated steatohepatitis (MASH), MASLD can predispose individuals to cirrhosis and hepatocellular carcinoma. Here, we characterized the first knockout mouse model for mitogen-activated protein kinase 15 (MAPK15) and revealed its critical role in controlling lipid homeostasis in the liver. Indeed, *Mapk15^-/-^* mice exhibited a MASLD-like phenotype, and hepatocellular models allowed us to demonstrate that dysregulated accumulation of lipids was due to increased expression and membrane localization of the CD36 fatty acid translocase. Consistently, *Mapk15^-/-^* mice exhibited elevated hepatic levels of CD36 and feeding them with a western-type diet significantly accelerated their progression to a MASH-like phenotype. Ultimately, transcriptomic analysis of human cohorts revealed increased liver expression of *MAPK15* in MASLD patients, compared to unaffected individuals, ultimately supporting a protective role for MAPK15 against this disease. Overall, our data highlight a critical role for MAPK15 in liver physiopathology, by contributing to maintain physiological intracellular levels of lipids in this tissue.

## INTRODUCTION

Lipid metabolism maintains a continuous state of dynamic equilibrium, regulated by dietary intake, de-novo lipogenesis (DNL), mobilization from adipose tissue, and the consumption of lipids, to ultimately meet body energy demands (Houten *et al*., 2016; Jeon *et al*., 2023; Samovski *et al*., 2023b). Disruption of the balance within these pathways that contribute to lipid homeostasis, through increased FAs uptake or enhanced DNL, are potential primary causes of excessive lipid storage, leading to significant deleterious effect and the development of important human diseases (Ding *et al*., 2017). Among them, metabolic dysfunction-associated steatotic liver disease (MASLD), formerly known as metabolic dysfunction-associated fatty liver disease (MAFLD) or non-alcoholic fatty liver disease (NAFLD) (Eslam *et al*., 2020; Rinella *et al*., 2023), has become the most common liver disorder worldwide and represents a leading cause of cirrhosis and hepatocellular carcinoma (HCC) (European *et al*., 2024). Indeed, MASLD occurrence is continuously rising, primarily due to poor dietary habits, with current estimation indicating a global prevalence exceeding 30% (European *et al*., 2024). Importantly, men exhibit a higher incidence and prevalence of MASLD compared to pre-menopausal women but not in relation to post-menopausal woman (Smiriglia *et al*., 2023). This suggests that female sex hormones may play a protective role in the pathogenesis of the disease (Smiriglia *et al*., 2023).

Primary criteria for defining MASLD include the presence of hepatic steatosis, characterized by the accumulation of triglycerides (TGs) in hepatocytes, and the absence of secondary causes such as excessive alcohol consumption, use of steatogenic drugs, or inborn metabolic disorders (European *et al*., 2024). MASLD encompasses a spectrum of liver conditions, ranging from simple liver steatosis (metabolic dysfunction-associated steatotic liver, MASL), to more severe forms such as metabolic dysfunction-associated steatohepatitis (MASH), progressing further to fibrosis and cirrhosis. This disease spectrum is often accompanied by increasing rate of obesity, metabolic syndrome, and type 2 diabetes mellitus (T2D) (European *et al*., 2024). Unfortunately, Resmetirom, an orally active, liver-directed, thyroid hormone receptor agonist, is the only drug for the treatment of MASH that has obtained positive results in a phase III clinical trial. Indeed, most pharmacological therapies for MASLD patients are currently directed to the management of cardiometabolic comorbidities (e.g., T2D and obesity) (European *et al*., 2024).

Lipid droplets (LDs) are the specialized organelles used by cells across a wide range of organism, from yeast to mammals, for storage of neutral lipids (Olzmann and Carvalho, 2019). The number, size and FA composition of LDs might differ reflecting the metabolic state of the cells. However, despite their heterogeneity, LDs share the characteristic of being surrounded by a phospholipid monolayer, intersperse by perilipins, to separate the hydrophobic neutral lipid core, mostly composed of triacylglycerol (TAG) and sterol esters, from the cytosol (Tauchi-Sato *et al*., 2002). LDs are formed from the membrane of the endoplasmic reticulum (ER), originating from the expansion of a neutral lipid lens between the membrane bilayer, which leads to LDs budding (Olzmann and Carvalho, 2019). Next, LDs can expand through droplet-droplet fusion, the transfer of TG from the ER membrane to the LDs, or through the direct synthesis of TG on the LDs surface (Olzmann and Carvalho, 2019). Conversely, FA can be mobilized from LDs either by lipolysis or by lipophagy, particularly during various phases of a cell lifespan or when subjected to stress (e.g., starvation) (Olzmann and Carvalho, 2019). Importantly, excessive FAs accumulation in LDs, when occurring chronically, may ultimately result in liver steatosis and obesity (Ding *et al*., 2017).

Mitogen-activated protein kinase 15 (MAPK15; ERK8; ERK7) is an atypical MAP kinase implicated in several cellular processes, such as cell proliferation (Iavarone *et al*., 2006; Xu *et al*., 2010; Colecchia *et al*., 2015; Jin *et al*., 2015; Rossi *et al*., 2016), genomic integrity (Groehler and Lannigan, 2010; Cerone *et al*., 2011; Rossi *et al*., 2016), secretion (Zacharogianni *et al*., 2011; Hasygar and Hietakangas, 2014), autophagy and mitophagy (Colecchia *et al*., 2012, 2015; Zhang *et al*., 2021; Franci *et al*., 2022), oxidative stress, aging, and cellular senescence (Franci *et al*., 2022, 2024). Interestingly, few studies have also already suggested a role for *MAPK15* in human diseases related to lipid metabolism. Among them, a genome-wide association study identified a polymorphism in this gene as a possible risk allele for childhood obesity (https://www.ebi.ac.uk/gwas/genes/MAPK15) (Comuzzie *et al*., 2012) while, more recently, *MAPK15* has been found associated to obesity-related traits (Ahn *et al*., 2019), consolidating evidence for its contribution to local fat distribution and obesity phenotypes. Ultimately, MAPK15 has been already described as an anti-anabolic kinase able to inhibit lipid storage in the fat body of Drosophila (Hasygar *et al*., 2021) and its expression to be stimulated in mouse livers upon high-fat diet (HFD)-induced MASLD (Wang *et al*., 2022).

Here, we characterized the first *Mapk15* knockout (KO, *Mapk15^-/-^*) mouse model and demonstrated a key role for this gene in controlling the accumulation of lipids in mouse hepatocytes, ultimately contributing to the establishment of MASLD. This status was associated with an overweight/obesity phenotype, insulin resistance and dyslipidemia. Importantly, we demonstrated that gene deletion in mice or downregulation in cultured cells led to increased expression and membrane localization of Cluster of Differentiation 36 (CD36, also known as SR-B2 or FAT) in hepatocytes. Ultimately, by feeding *Mapk15^-/-^*mice with a lipid-and carbohydrate-rich western-style diet (WD), we demonstrated that the loss of this gene greatly accelerated the progression of MASLD towards more advanced pathological features of MASH, such as hepatocytes damage, liver inflammation and fibrosis.

## RESULTS

### *Mapk15* gene deletion induces MASLD

To characterize the specific phenotypes of the newly generated *Mapk15* KO (*Mapk15^-/-^*) mouse model in the C57BL/6J genetic background (**Fig. S1A** and **Fig. S1B**), we initiated our analysis by examining its baseline morphometric traits and blood chemistry profile. Upon feeding with a standard diet (SD), male *Mapk15^-/-^* mice showed a significantly higher body weight (BW) when compared to age-and sex-matched wild type (*Mapk15^+/+^*, WT) mice (**Fig. 1A**). Indeed, male *Mapk15^-/-^*mice showed an “overweight” phenotype, as demonstrated by visual examination (**Fig. 1B**), paralleled by a significantly higher BCS when 24-weeks old (Ullman-Culleré and Foltz, 1999) (**Fig. S2**), and by the evidence of more abundant visceral adipose deposits upon necroscopic analysis (**Fig. 1C**) with a significant increased volume of the single adipocytes, demonstrated by histologic analysis (**Fig. 1D**). Twenty-four-weeks old male *Mapk15^-/-^* and *Mapk15^+/+^*mice showed equivalent fasting serum levels of TGs and transaminases (alanine aminotransferase, ALT, and aspartate aminotransferase, AST), but significantly higher values of fasting serum cholesterol in *Mapk15^-/-^*animals (**Fig. 1E**). Male *Mapk15^-/-^* mice also showed significantly higher fasting serum levels of glucose and insulin than *Mapk15^+/+^* animals, demonstrating increased insulin resistance index, evaluated by the homeostatic model assessment for insulin resistance (HOMA IR) method (Matthews *et al*., 1985; Fraulob *et al*., 2010) (**Fig. 1F**). Overall, our data therefore supported the existence of important metabolic dysfunctions (“metabolic syndrome”) (Kanwar and Kowdley, 2016) resulting from the deletion of the *Mapk15* gene.

**Figure 1.**
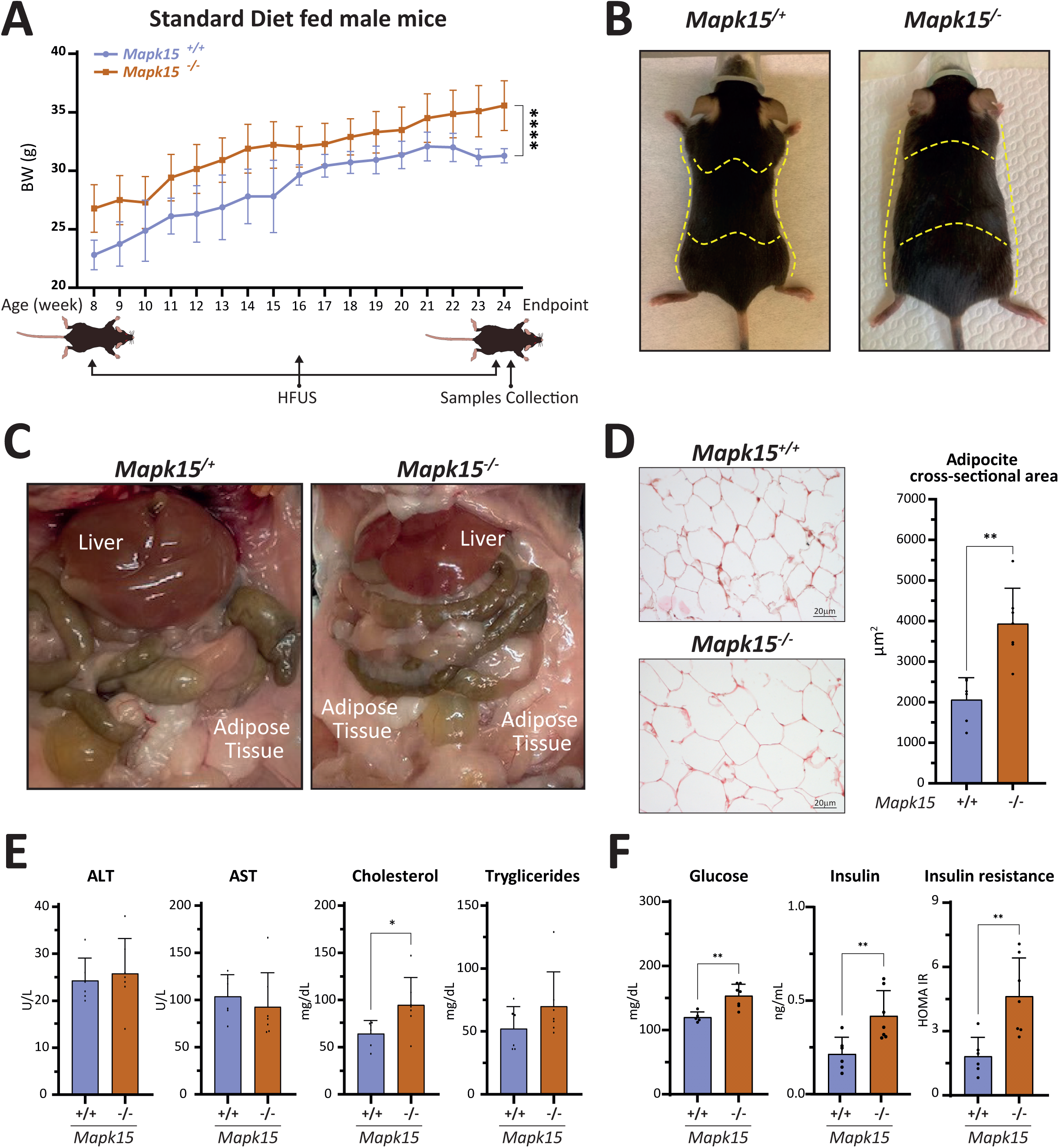
– *Mapk15^-/-^* mice show metabolic syndrome. **A** *Mapk15^+/+^* and *Mapk15^-/-^* mice were fed with a standard diet (SD) and data were collected from 8 to 24 weeks of age. Mice were weighed every week, visceral ultrasound data were acquired at 8, 16 and 24 weeks of age while blood samples were collected at the end of the study. Two-way ANOVA-Mixed effect model -Repeated measures, interaction age x genotype: ****p ≤ 0.0001. **B** Representative images of the appearance of *Mapk15^+/+^* and *Mapk15^-/-^* mice at 24 weeks of age. **C** Necroscopic analysis of *Mapk15^+/+^* and *Mapk15^-/-^* SD-fed mice at 24 weeks of age. **D** Histologic examination of visceral adipose tissue from *Mapk15^+/+^*and *Mapk15^-/-^* SD-fed mice at 24 weeks of age. Adipocyte cross-sectional area was quantified and reported in the accompanying histogram (µm^2^). **E** Blood samples from 24-week-old *Mapk15^+/+^* and *MAPK15^-/-^*SD-fed mice were collected and analyzed for indicated markers: ALT, AST, cholesterol and TG. **F** Same as in **E** but analyzing glucose and insulin. Insulin resistance values (HOMA IR) were also calculated. One way ANOVA – Mann Whitney non-parametric test: *p ≤ 0.05; **p ≤ 0.01.

Ultrasound analysis of SD-fed male *Mapk15^-/-^* mice, compared to their *Mapk15^+/+^* counterparts, also showed, at different time-points, *in vivo* changes of hepatic features indicative of mild parenchymal steatotic alterations (**Fig. 2A**). Indeed, 24-weeks old SD-fed *Mapk15^-/-^* male mice showed a diffuse increase of parenchymal echogenicity and echotexture heterogeneity, and frequent (43%) incidence of isoechoic appearance of the caudate liver lobe compared with the right kidney cortex (**Fig. 2A**), although differences in parametric values such as hepatorenal index (HRI) and liver echogenicity did not reach statistical significance (**Fig. S3**).Therefore, to definitely ascertain the presence of hepatic accumulation of lipids in *Mapk15^-/-^*male mice, these were sacrificed when 24-weeks old, and livers, which appeared paler although of similar weights when compared to *Mapk15^+/+^* mice (**Fig. 2B**), were subjected to histological analysis with a specific fat-soluble dye for intracellular lipid stain (i.e., Oil Red O), ultimately revealing widespread hepatocellular lipid accumulation (Benedict and Zhang, 2017) (**Fig. 2C**). Therefore, our findings of overweight phenotype, insulin resistance and dyslipidemia in presence of hepatic accumulation of lipids strongly suggested that deletion of the *Mapk15* gene promoted the onset of a MASLD-like phenotype in mice (European *et al*., 2024). Importantly, the histological evaluation of livers from male *Mapk15^-/-^* mice did not reveal evident accumulation of inflammatory cells (**Fig. 2D**) and/or liver fibrosis (**Fig. 2E**), thus excluding a more advanced liver disease (Benedict and Zhang, 2017). To investigate the potential role of sex in the development of MASLD (Smiriglia *et al*., 2023), we performed equivalent studies in female mice. Interestingly, while female *Mapk15^-/-^* mice also showed an increased tendency to accumulate lipids in their livers, this phenotype was milder than in their male counterparts, as demonstrated by *ex vivo* non-significant differences in the histological appearance of livers between *Mapk15^-/-^* and *Mapk15^+/+^*mice at Oil Red O (**Fig. S4A**) and H&E (**Fig. S4B**) staining. Therefore, our data confirm available literature showing that female mice have a reduced tendency to accumulate hepatic TGs and develop steatosis (Norheim *et al*., 2017; Smiriglia *et al*., 2023), recapitulating the observation of a higher prevalence of MASLD in men when compared to pre-menopausal women (but not to those in post-menopausal status) (Smiriglia *et al*., 2023).

**Figure 2.**
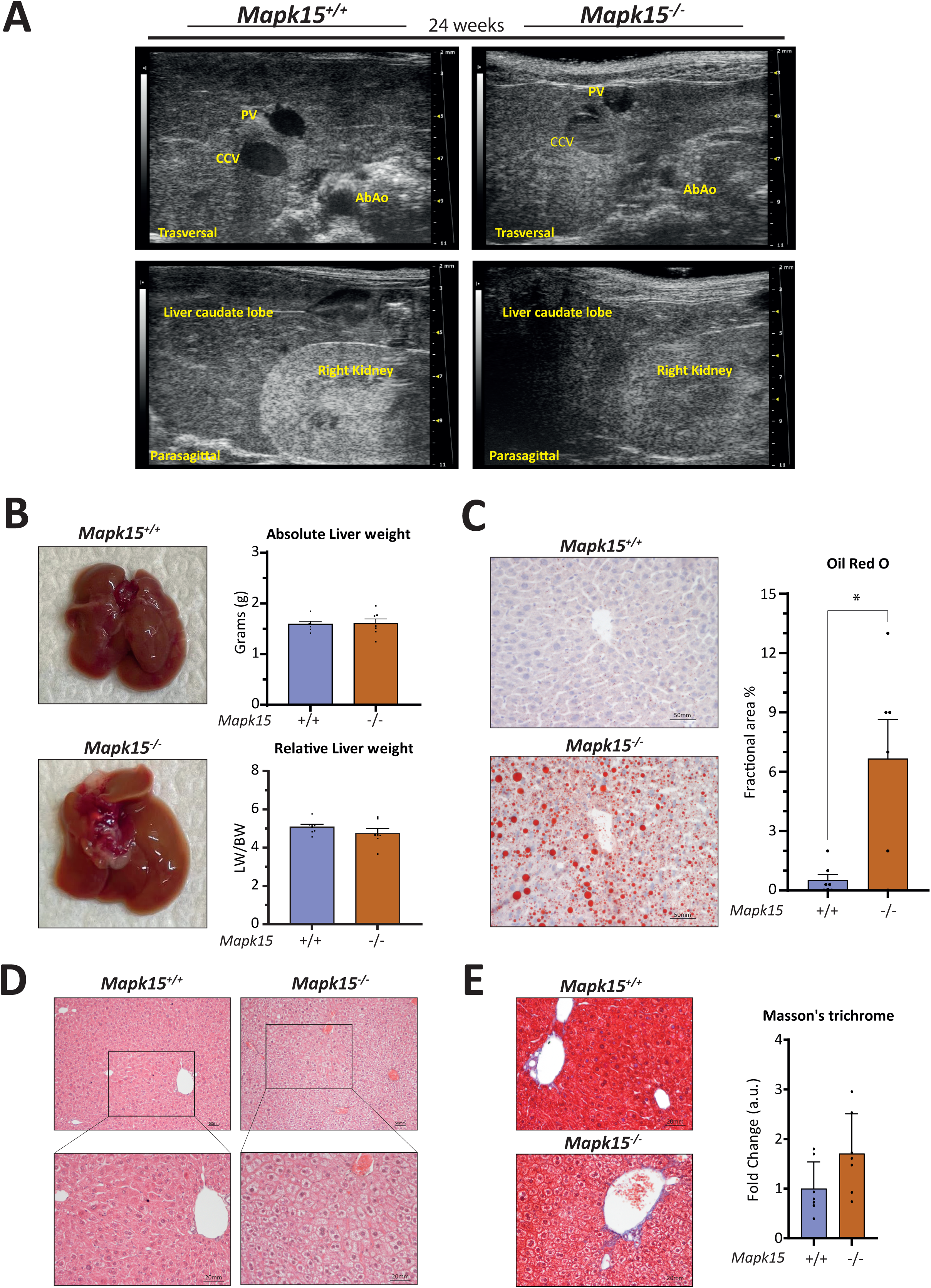
–*Mapk15^-/-^* mice show hepatic accumulation of lipids. **A** Representative images of ultrasound acquisition at 24 weeks of age. **B** Representative gross liver pictures and absolute (liver weight, LW) and relative (body weight/liver weight, BW/LW) liver weights. **C** Representative Oil Red O staining of histological samples from livers of *Mapk15^+/+^* and *Mapk15^-/-^* male mice. The accompanying graph scores fractional area of Oil Red O staining. **D** Representative haematoxylin and eosin staining of histological samples from *Mapk15^+/+^*and *Mapk15^-/-^* SD-fed male mice livers. **E** Representative Masson’s trichrome staining of histological sections from *Mapk15^+/+^* and *Mapk15^-/-^* SD-fed mice livers. The accompanying graph scores the blue collagen fibres as fold changes comparing WT (n=7) and KO (n=7) liver. One way ANOVA – Mann Whitney nonparametric test: *p ≤ 0.05.

### *MAPK15* depletion stimulates accumulation of lipid droplets in hepatocellular in vitro models

Given the observed increase in intracellular lipid accumulation in the liver of *Mapk15^-/-^* mice, we next decided to use the HepG2 hepatoblastoma cell line as a model to further investigate the molecular mechanisms underlying MAPK15-dependent regulation of lipid metabolism, with the aim of uncovering key pathways contributing to this phenotype. As LDs are the most important site for intracellular storage of lipids (Olzmann and Carvalho, 2019), we used the BODIPY 493/503 probe to evaluate the amount of these organelles in HepG2 cells transiently depleted for *MAPK15* expression by a classical RNA interference approach. Indeed, HepG2 cells transfected with siRNA against *MAPK15* (siMAPK15) (**Fig. S5**) showed a significant increase in LD amounts, when analysed by confocal microscopy, compared to control siRNA-transfected cells (siSCR) (**Fig. 3A**). This result was also confirmed by flow cytometry analysis quantifying LDs mean fluorescence intensity, using two different *MAPK15* specific siRNAs (**Fig. 3B**). Importantly, to confirm that such mechanism was conserved also in different cellular models, we also used hepatoma HepaRG cell line interfered for *MAPK15* expression (**Fig. S6A**) and demonstrated also in these cells an increase in the amounts of LDs both by confocal microscopy (**Fig. S6B**) and cytofluorimetric (**Fig. S6C**) approaches. As MASLD induction can be modelled in HepG2 cells *in vitro* by short term treatments with palmitic acid (PA) and/or oleic acid (OA) (Smiriglia *et al*., 2023), LDs content was also examined by confocal microscopy (**Fig. 3C-D**) and flow cytometry analysis (**Fig. 3E**) after 24 h treatment with these FAs, demonstrating also in these conditions an increase in number and size of LDs in siMAPK15 cells when compared to siSCR cells. Importantly, this result also suggested that an increased uptake of exogenous FAs could be responsible for the enhanced accumulation of lipids inside hepatic cells. Indeed, we demonstrated an increased cellular uptake of exogenously added fluorescent palmitate (BODIPY-C16) in siMAPK15 HepG2 cells, when compared to their siSCR counterpart (**Fig. 3F**).

**Figure 3.**
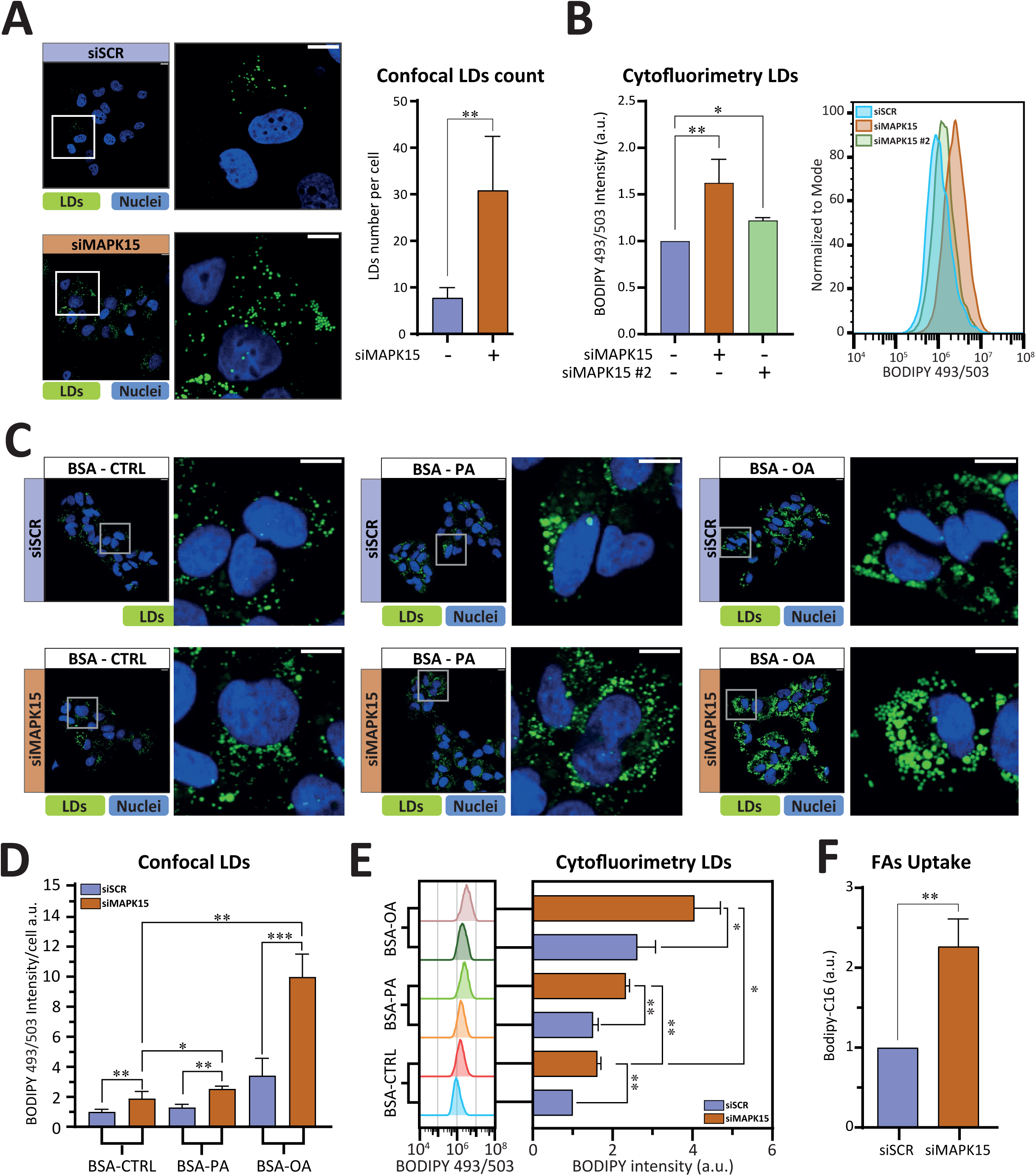
– *MAPK15* depletion increases LD content in hepatocellular model systems. **A** Representative confocal microscopy images showing lipid droplets (LDs) content in *MAPK15*-interfered (72 h) HepG2 cells (siMAPK15), stained with BODIPY 493/503. The accompanied graph shows quantification of LDs from 10 representative microscopy fields signal per cell ± SD, using the quantitation Module of the Volocity software. **B** Same as in **A** but using flow cytometry analysis to score cellular content of LDs fluorescence. **C** Representative confocal microscopy images showing lipid droplets after 24 h treatment with 200 µM of FAs (BSA-Palmitic acid and BSA-Oleic Acid; BSA as control), in *MAPK15*-interfered (72 h) cells and relative controls. **D** Densitometric analysis of **C**. BODIPY 493/503 fluorescence was scored from five representative microscopy fields signal per cell ± SD. **E** Same as in **C** but using flow cytometry analysis to score LDs cellular content fluorescence after treatments with FAs. **F** Cytofluorometric quantification of exogenous FAs uptake using BODIPY-C16 (2 µM). Scalebar 10 µm. Unpaired t-test with Welch’s correction: *p ≤ 0.05, **p ≤ 0.01, ***p ≤ 0.001, ****p ≤ 0.0001.

### Accumulation of LDs in MAPK15 depleted cells depends on the induction of CD36 expression and membrane localization

Approximately 59% of hepatic fat originates from circulating free FAs, while *de novo* lipogenesis accounts for about 26%, and dietary sources for the remaining 15% (Cohen *et al*., 2011). Several FAs transport proteins, particularly FATP1-6 (SLC27 protein family) and CD36, have been implicated in mediating the transport of exogenous long and very-long chain fatty acids (LCFA and VLCFA) through the cellular membrane (Samovski *et al*., 2023a) (**Fig. 4A**). Importantly, it is well known that CD36 increases FAs uptake, and, in the liver, drives hepato-steatosis onset and might contribute to its progression to MASH (Rada *et al*., 2020). Therefore, we hypothesized that the increase of LDs content observed in siMAPK15 HepG2 cells may be attributed to enhanced uptake of exogenous free FA, potentially mediated by CD36. Indeed, upon downregulation of *MAPK15*, HepG2 cells showed increased expression of CD36 at both mRNA (**Fig. 4B**) and protein (**Fig. 4C**) level.

**Figure 4.**
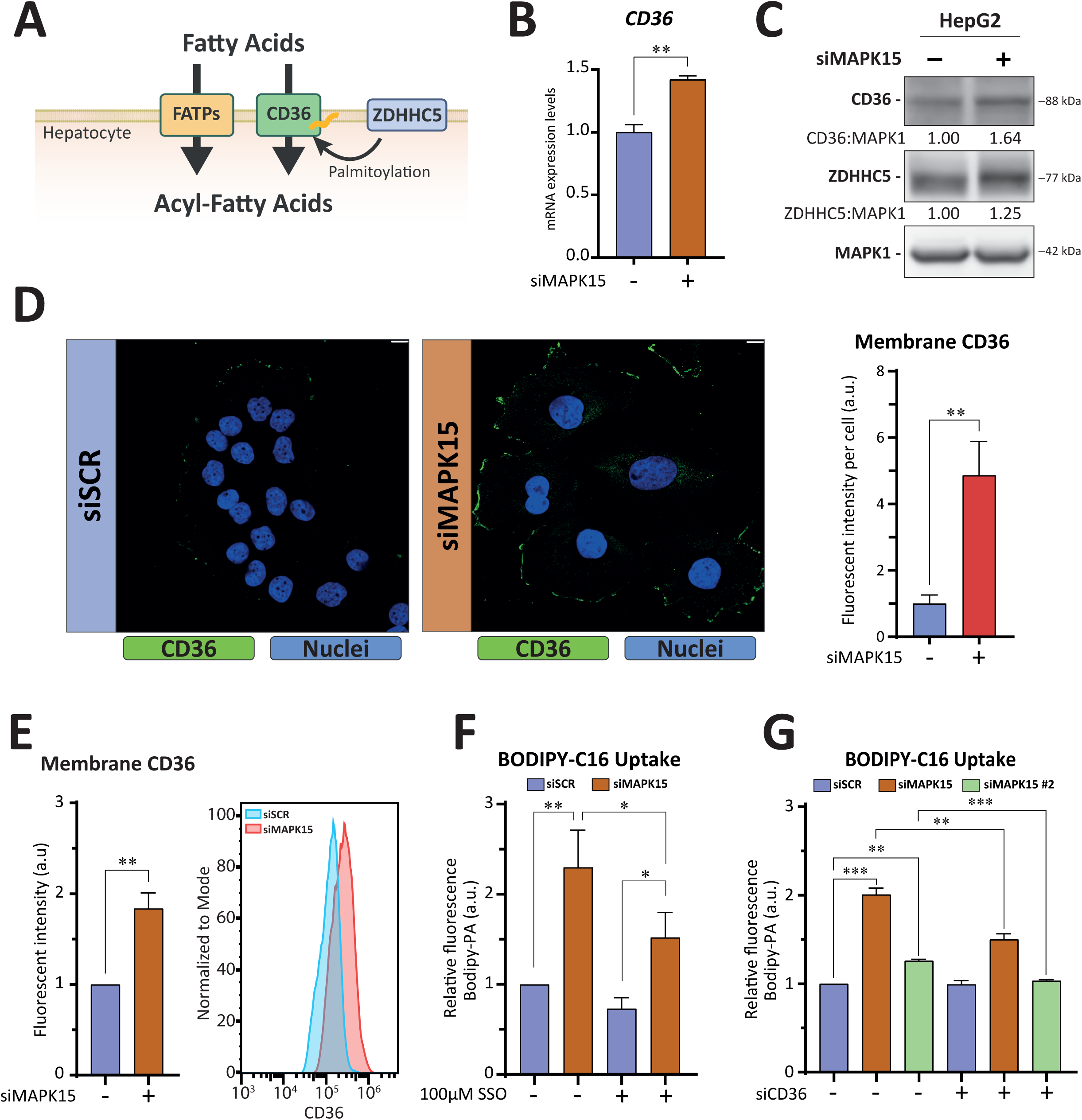
– *MAPK15* depletion increases CD36 expression and membrane localization. **A** A cartoon illustrating the transport of exogenous long-chain and very-long-chain fatty acids across the cellular membrane. **B** Quantitative reverse transcription polymerase chain reaction (RT-qPCR), to monitor mRNA expression of *CD36* from HepG2 cells interfered for *MAPK15* expression (72 h). **C** Representative immunoblots of CD36 and ZDHHC5 protein expression from HepG2 cells interfered for *MAPK15* expression (72 h). **D** Representative confocal microscopy images of membrane-localized CD36 from HepG2 cells interfered for *MAPK15* expression (72 h). Intensitometric analysis of CD36 fluorescence from five representative microscopy fields signal per cell ± SD is shown in the accompanying histogram. **E** Cytofluorometric quantification of the membrane-localized CD36 protein, in HepG2 cells interfered for *MAPK15* expression (72 h). **F** Cytofluorometric quantification of BODIPY-C16 (2 µM) in HepG2 cells interfered for *MAPK15* expression (72 h) and treated with 100 µM SSO in the absence of FBS for 6 h. **G** Same as in **F** but treating *MAPK15* interfered HepG2 cells with a specific siRNA against *CD36*. Scalebar 10 µm. Unpaired t-test with Welch’s correction: *p ≤ 0.05, **p ≤ 0.01, ***p ≤ 0.001.

Subcellular localization of CD36 at the plasma membrane contributes to determining its efficacy in mediating free FAs uptake. Crucially, CD36 palmitoylation mediated by ZDHHC5 is required for its plasma membrane localization (Wang *et al*., 2019). As we also observed increased ZDHHC5 protein levels in MAPK15 downregulated cells (**Fig. 4C**), we next evaluated CD36 membrane localization in these experimental conditions. Indeed, increased CD36 membrane localization was demonstrated in HepG2 cells interfered for *MAPK15* expression, when scored by both confocal microscopy (**Fig. 4D**) and by flow cytometry (**Fig. 4E**). Next, to demonstrate a causal role for CD36 in the observed increase of LDs in MAPK15-interfered cells, we used a specific CD36 inhibitor, Sulfo-N-succinimidyl oleate (SSO) (Kuda *et al*., 2013). Indeed, SSO significantly rescued palmitate uptake induced by *MAPK15* downregulation, in HepG2 cells (**Fig. 4F**). CD36 mRNA and protein expression were also similarly increased by downregulating MAPK15 expression in HepaRG cells (**Fig S7A**), and SSO prevented the increase of palmitate uptake induced by *MAPK15* downregulation also in HepaRG cells (**Fig. S6B**). As an additional confirmation of the SSO pharmacological rescue approach, we also interfered the expression of *CD36* in HepG2 cells by specific siRNA (siCD36) to rescue the effect induced by MAPK15 downregulation (**Fig. S8**). Indeed, we observed a reduction in the cellular uptake of palmitate also in these experimental conditions (**Fig. 4G**), overall indicating that the observed increase in the expression of *CD36* is necessary for enhanced accumulation of exogenous FAs in siMAPK15 HepG2 cells. Accordingly, mouse livers from *Mapk15^-/-^*mice also expressed increased amounts of *Cd36* mRNA as compared to livers from *Mapk15^+/+^* mice, while expression of other genes involved in transmembrane fatty acid transport (i.e., *Fatp1*, *Fatp2*, *Fatp5, Fatbp1* and *Fatb4*) were not significantly affected by *Mapk15* gene deletion (**Fig. 5A**). Livers from *Mapk15^-/-^* mice also expressed higher protein levels of both CD36 and ZDHHC5 (**Fig. 5B**) and immunohistochemistry (IHC) analysis showed increased CD36 localization at the plasma membrane (**Fig. 5C**), overall supporting an *in vivo* role for CD36 in the lipid metabolic rewiring governed by MAPK15 in liver cells.

**Figure 5.**
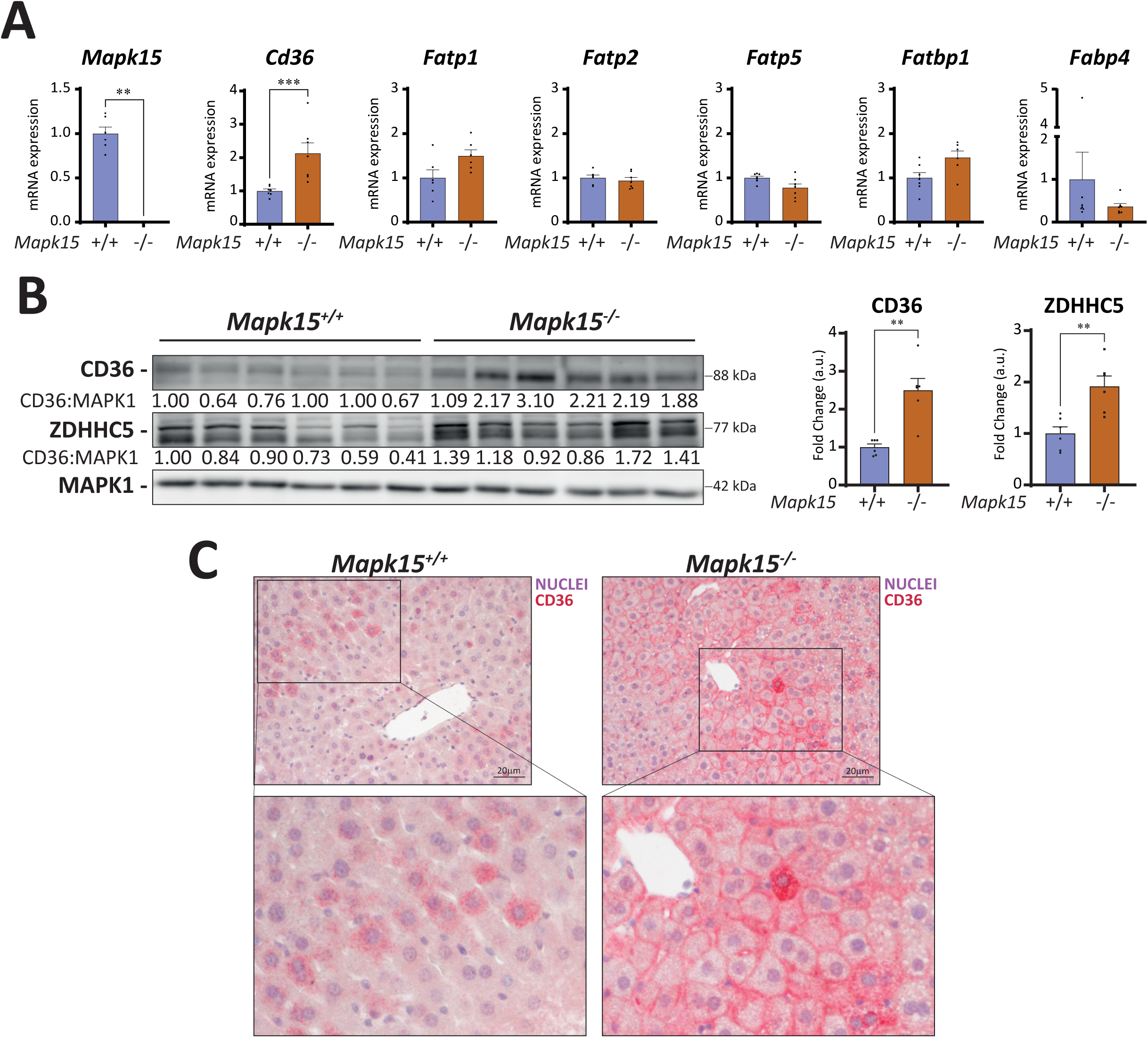
– *Mapk15^-/-^* mice show increased *CD36* expression and membrane localization. **A** RT-qPCR was performed on livers from *Mapk15^+/+^* and *Mapk15^-/-^* mice to evaluate mRNA levels of indicated genes (*Mapk15, Cd36, Fatbp1, Fatbp4, Fatp1, Fatp2, and Fatp5*). **B** Representatives immunoblots of mice liver protein extracts for indicated proteins (CD36, ZDHHC5 and MAPK1). Densitometric analysis of average protein expression levels in livers from *Mapk15^+/+^* and *Mapk15^-/-^*mice is shown in the accompanying graph. **C** IHC analysis of CD36 in liver sections of both genotypes. One way ANOVA – Mann Whitney non-parametric test: *p ≤ 0.05, **p ≤ 0.01, ***p ≤ 0.001.

### DNL does not contribute to intracellular accumulation of lipids in MAPK15 depleted/deleted cells

DNL is largely controlled at the transcriptional levels by two master transcription factors, i.e. sterol regulatory element binding protein 1 (SREBP-1) and carbohydrate-responsive element-binding protein (ChREBP), which stringently regulate the expression of key rate-limiting lipogenic enzymes (e.g., Acetyl-CoA Carboxylase 1, ACC; Fatty Acid Synthase, FASN; Stearoyl-CoA Desaturase-1, SCD1) (Jeon *et al*., 2023; Esler and Cohen, 2024) (**Fig. 6A**). Importantly, together with the uptake of exogenous FAs, DNL represents another key mechanism driving hepatic FAs accumulation. Indeed, hepatic steatosis typically occurs when the rate of one or both these processes exceed the capacity of the liver to oxidize FAs or secrete TGs via very-low-density lipoproteins (VLDL) (Esler and Cohen, 2024). Still, *MAPK15*-interfered HepG2 cells showed significantly reduced lipid biosynthesis when pulsed with radiolabelled ^14^C-glucose (**Fig. 6B**), suggesting that DNL did not contribute to the observed intracellular accumulation of lipids observed when reducing *MAPK15* expression. Accordingly, we next observed that acute downregulation of *MAPK15* by RNA interference, in HepG2 cells, strongly reduced the expression of *SREBP-1* (**Fig. 6C**) and its specific transactivating function, measured through the transfection of a reporter gene under the control of the sterol regulatory element (SRE) (Dooley *et al*., 1998) (**Fig. 6D**). Similarly, also the expression of the *ChREBP* transcription factor, which acts complementarily to SREBP-1 to induce lipogenesis (Jeon *et al*., 2023), was significantly reduced in MAPK15-interfered HepG2 cells (**Fig. 6E**). Consequently, mRNA (**Fig. 6F**) and protein (**Fig. 6G**) levels of SREBP-1 and ChREBP transcriptional targets, i.e., *ACC*, *FASN* and *SCD1*, were also reduced, in MAPK15-interfered HepG2 cells. Interestingly, the expression of *Srebp-1* and *Chrebp* (**Fig. 6H**), and of their target genes (**Fig. 6I**) did not significantly change in the livers of *Mapk15^-/-^* mice, as compared to those from *Mapk15^+/+^* mice, overall demonstrating that DNL does not positively contribute to the lipid accumulation observed in MAPK15-depleted hepatic cells.

**Figure 6.**
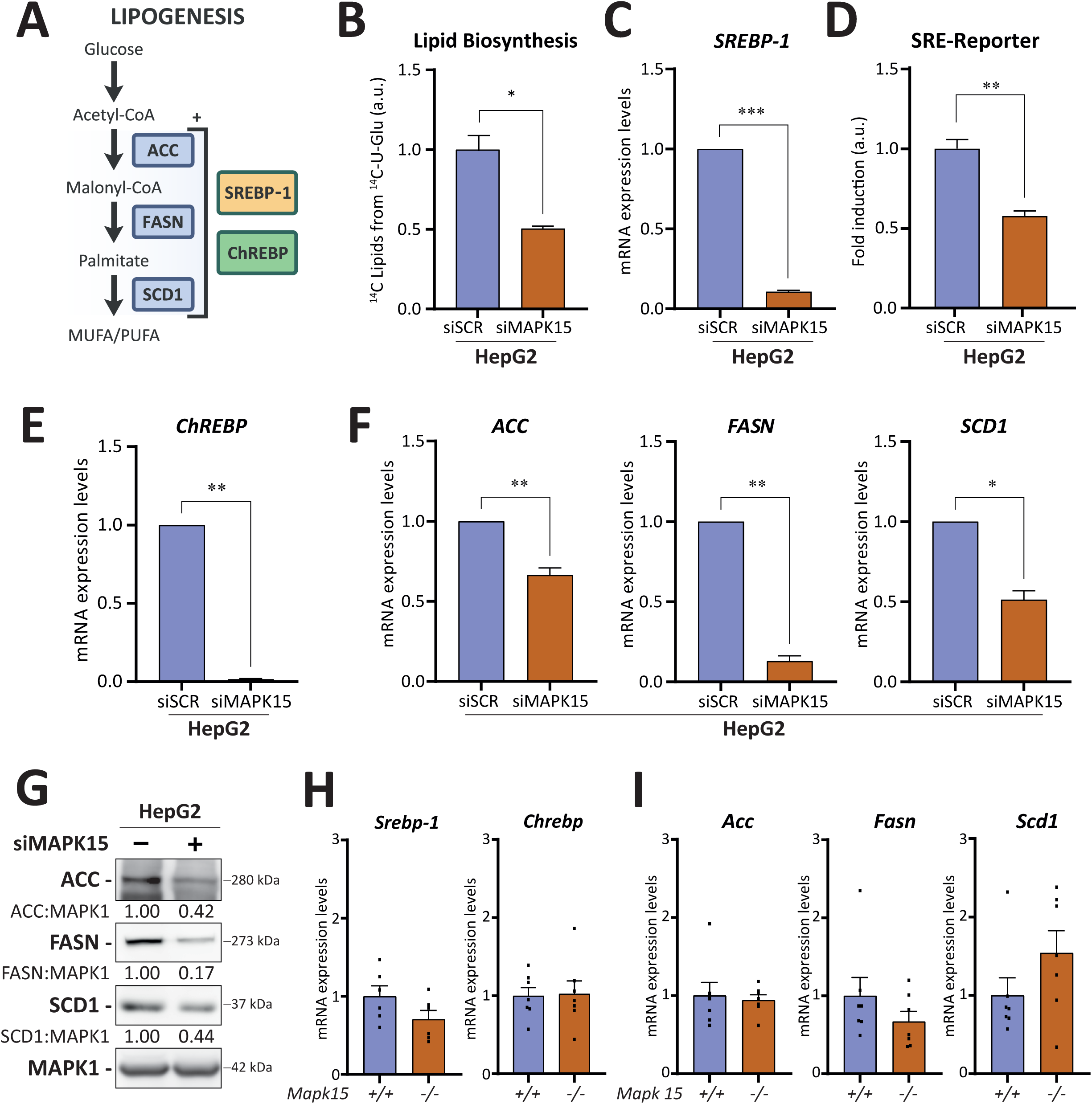
– *De novo* lipogenesis does not contribute to LD accumulation in *MAPK15* depleted cells. **A** Cartoon illustrating the main pathway for *de novo* lipogenesis. **B** Functional assay to evaluate lipid biosynthesis from ^14^C-U-labeled glucose, in MAPK15-interfered (72 h) HepG2 cells. Lipids were extracted, and the radioactive signal was measured to track metabolite incorporation into lipids. Each value was normalized to cell counts. **C** RT-qPCR evaluation of *SREBP-1* mRNA level in MAPK15-interfered (72 h) HepG2 cells. **D** Activation of a sterol regulatory element (SRE) luciferase reporter plasmid transfected in MAPK15-interfered (72 h) HepG2 cells. **E** Evaluation of ChREBP mRNA expression in MAPK15-interfered (72 h) HepG2 cells. **F** RT-qPCR performed to evaluate mRNA levels of *ACC*, *FASN* and *SCD1* upon *MAPK15* RNA interference of HepG2 cells. **G** Representative immunoblots of MAPK15-interfered (24 h) HepG2 cell extracts to evaluate expression of ACC, FASN, SCD1 proteins. **H** RT-qPCR was performed on livers from *Mapk15^+/+^* and *Mapk15^-/-^*mice to evaluate mRNA levels of the *Srebp-1* and *Chrebp* genes. The graphs show average expression of the different genes in the livers of *Mapk15^+/+^*and *Mapk15^-/-^* mice. **I** Same as in **H** but evaluating *Acc*, *Fasn* and *Scd1* gene expression. Unpaired t test with Welch’s correction: *p ≤ 0.05, **p ≤ 0.01, ***p ≤ 0.001.

### Western-type diet synergizes with loss of MAPK15 to induce MASH

To positively define a role for this gene in protecting mammalian liver from steatogenic insults, we next tested a potentially synergizing effects of a high-fat, western-style diet (WD) on the livers of *Mapk15^-/-^*mice in accelerating progression towards the development of more severe pathological features of MASLD. Indeed, in humans, a diet high in unhealthy fats (such as saturated and trans-fats) is considered an independent risk factor for the development and progression of MASLD (Benedict and Zhang, 2017) and, correspondingly, high fat diet-fed mouse models have recently emerged as the most balanced in terms of metabolic, histologic and transcriptomic similarities to human disease (Vacca *et al*., 2024). Therefore, starting from the 8^th^ up to the 24^th^ week of age (total 16 weeks), male *Mapk15^+/+^* and *Mapk15^-/-^* mice were switched from the SD to a lipid-rich WD and analysed for different markers pathognomonic of human MASLD. WD-fed *Mapk15^-/-^* mice showed an average BW significantly higher than age-matched *Mapk15^-/-^* mice (**Fig. 7A**) and a clearly overweight/obese phenotype, as demonstrated by visual examination (**Fig. 7B**) and by a significantly higher BCS (**Fig. S9**). Visceral adipose deposits in *Mapk15^-/-^* mice were increased when compared to *Mapk15^+/+^*mice (**Fig. 7C**), as evidenced by enlarged volume of the associated adipocytes (**Fig. 7D**). Liver ultrasound analysis of male *Mapk15^-/-^* mice showed a higher incidence of more severe steatosis features both in the early and late experimental phases (**Fig. S10A**), and significantly higher HRI (**Fig. S10B**) and liver echogenicity (**Fig. S10C**) at the experimental endpoint, compared to *Mapk15^+/+^*mice. These findings were next confirmed by necropsy at 24 weeks of age, showing a paler appearance of livers from WD-fed *Mapk15^-/-^* mice (**Fig. 7E**), absolute liver weight (LW) and relative liver weight (body weight/LW, BW/LW) significantly higher in *Mapk15^-/-^* mice (**Fig. 7F**), and much larger cytoplasmic lipid accumulation in their hepatocytes (**Fig. 7G**). WD-fed *Mapk15^-/-^* mice also showed significantly higher baseline fasted serum levels of glucose and insulin than *Mapk15^+/+^*animals, with higher insulin resistance (HOMA IR) (**Fig. 7H**). Ultimately, WD-fed *Mapk15^-/-^*mice showed significantly higher baseline fasted serum values of cholesterol but a similar TG amount, and higher serum levels of ALT (**Fig. 8I**), a specific biomarker of hepatic alterations such as steatosis, due to its greater concentration in the liver compared with other tissues (Cui *et al*., 2018). Importantly, WD-fed *Mapk15^-/^*^-^ mice still expressed significantly increased levels and plasma membrane localization of CD36 protein compared to corresponding *Mapk15^+/+^* animals, as demonstrated by western blot (**Fig. 7L**) and IHC analysis (**Fig. 7M**), respectively.

**Figure 7.**
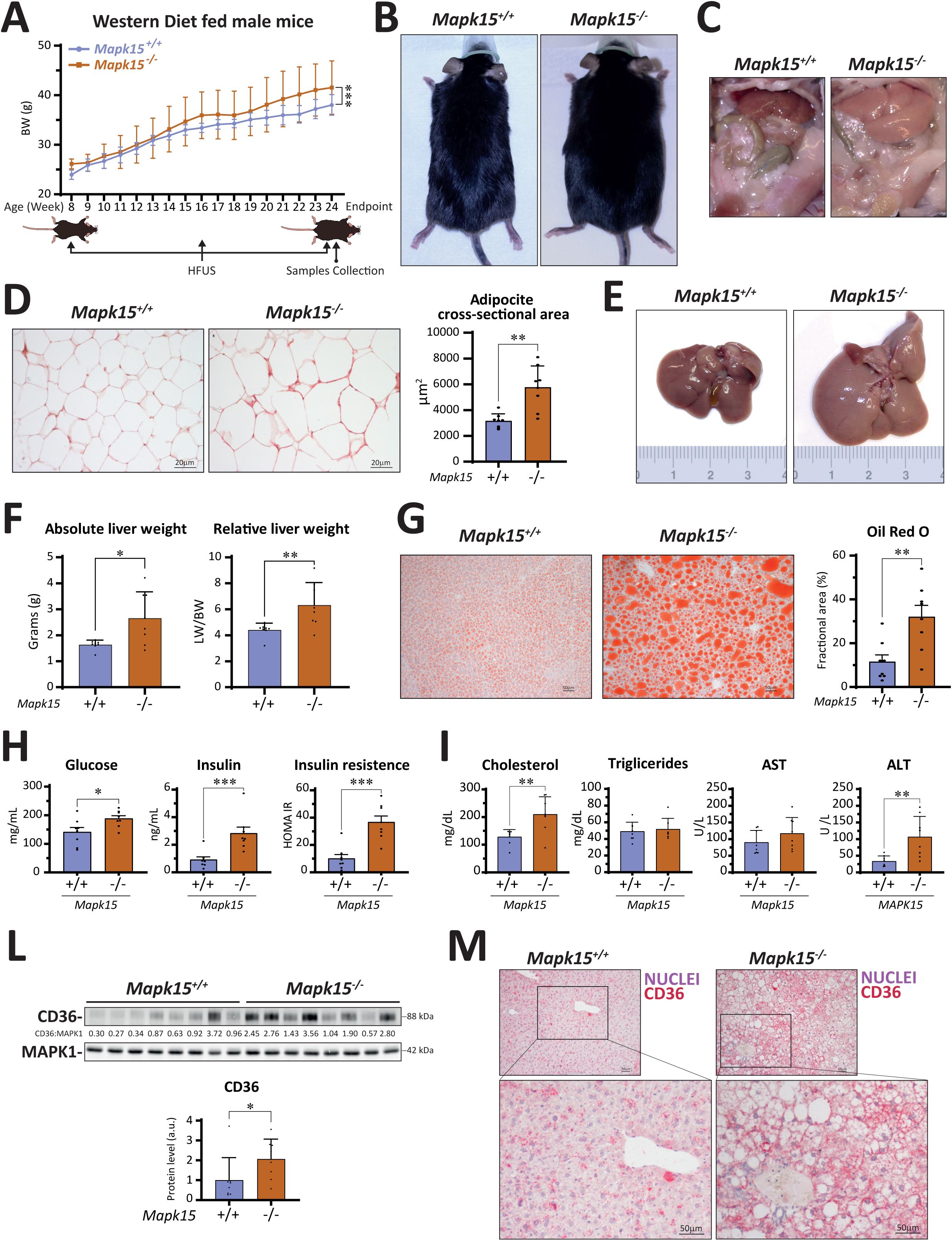
– *Mapk15* gene deletion cooperates with western-type diet to accelerate the progression of the MASLD phenotype. **A** From 8 to 24 weeks of age, *Mapk15^+/+^* and *Mapk15^-/-^* mice were fed with a wester-type diet (WD). Mice were weighed every week, as reported in the graph, and visceral ultrasound data were acquired at 8, 16 and 24 weeks of age. Blood samples were collected at the endpoint (24 weeks). Two-way ANOVA-Mixed effect model -Repeated measures, interaction age x genotype: ***p<0.001. **B** Representative images of the appearance of *Mapk15^+/+^* and *Mapk15^-/-^* WD-fed mice. **C** Necroscopic analysis of *Mapk15^+/+^* and *Mapk15^-/-^* WD-fed mice. **D** Histologic examination of visceral adipose tissue from *Mapk15^+/+^*and *Mapk15^-/-^* WD-fed mice. The accompanying graph shows the average adipocyte cross-sectional area from *Mapk15^+/+^* (n=8) and *Mapk15^-/-^* (n=8) mice. **E** Representative gross liver pictures from *Mapk15^+/+^* and *Mapk15^-/-^* WD-fed mice. **F** Absolute (liver weight, LW) and relative (liver weight/body weight, LW/BW) liver weights of *Mapk15^+/+^* and *Mapk15^-/-^* WD-fed mice. **G** Representative Oil Red O staining of histological samples from livers of *Mapk15^+/+^* and *Mapk15^-/-^* WD-fed mice. The accompanying graph scores the fractional area of Oil Red O staining. **H** Blood samples from 24-week-old *Mapk15^+/+^* and *Mapk15^-/-^*WD-fed mice were collected and analyzed for indicated markers (glucose and insulin). Insulin resistance values (HOMA IR) were also calculated. **I** Same as **H** but for indicated markers (cholesterol, triglycerides, AST and ALT). **L** Immunoblot analysis of CD36 protein expression in livers from *Mapk15^+/+^*and *Mapk15*^-/-^ WD-fed mice. Quantification of average expression in the different genotypes is also shown in the accompanying graph. **M** Representative IHC of CD36 in liver sections of both genotypes. One-way ANOVA – Mann Whitney nonparametric test: *p ≤ 0.05, **p ≤ 0.01, ***p ≤ 0.001.

**Figure 8.**
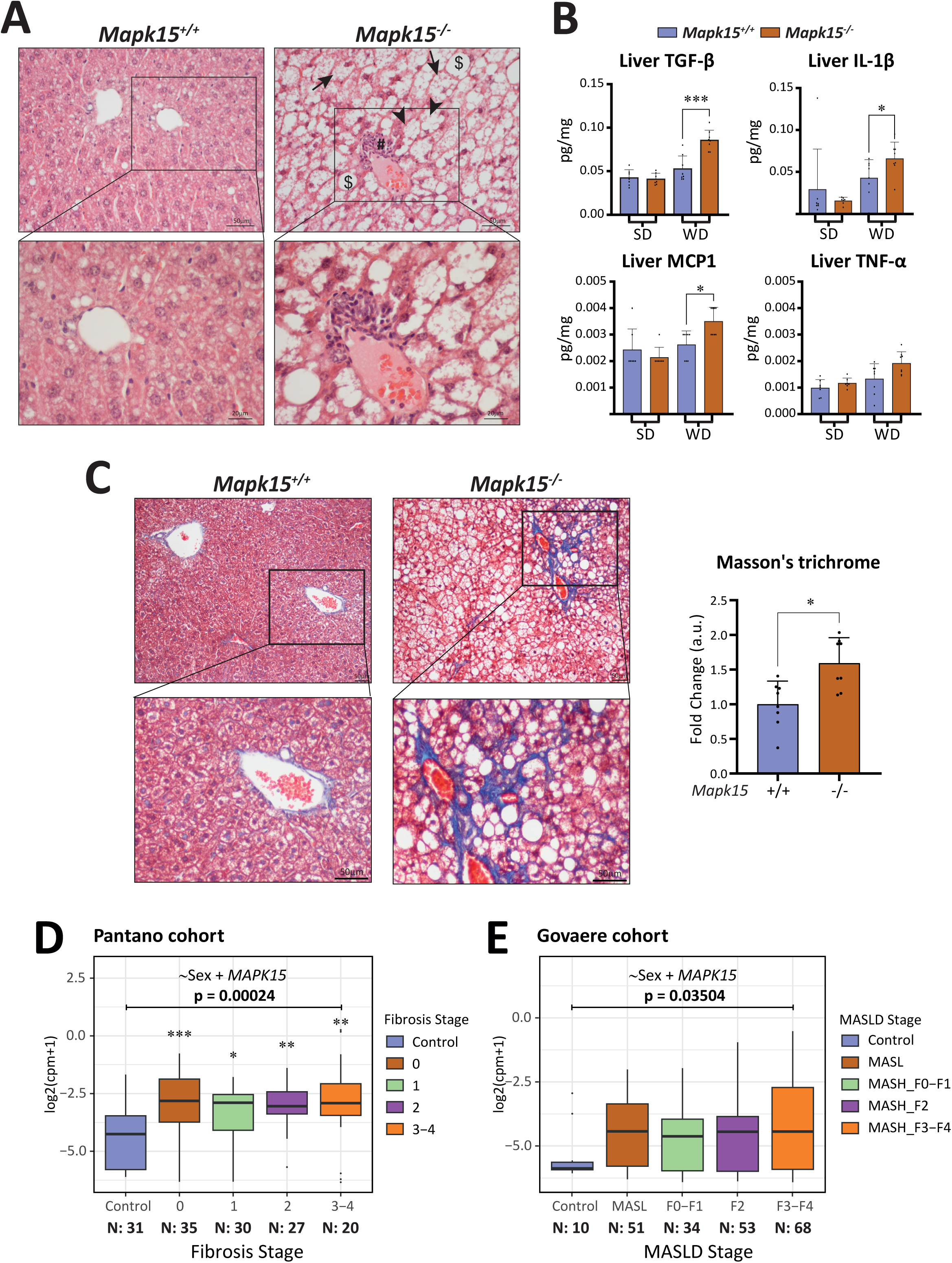
– Loss of *Mapk15* in combination with WD leads to hepatic damage and MASH development. **A** Representative haematoxylin and eosin staining on histological samples from livers of *Mapk15^+/+^*and *Mapk15^-/-^* WD-fed mice. #: inflammatory infiltrates; arrows: ballooning hepatocytes; arrowheads: micro-vesicular steatosis; $: macro-vesicular steatosis. **B** Local protein levels of indicated proinflammatory cytokines (TGF-β, IL-1β, MCP1, and TNF-α) from livers of *Mapk15^+/+^* and *Mapk15^-/-^*WD-fed mice. **C** Representative Masson’s trichrome staining of histological sections from *Mapk15^+/+^* and *Mapk15^-/-^* WD-fed mice. The accompanying graph scores the blue collagen fibers as fold changes comparing *Mapk15^+/+^* (n=8) and *Mapk15^-/-^* (n=8) liver. One-way ANOVA – Mann Whitney nonparametric test: (*p ≤ 0.05, **p ≤ 0.01, ***p ≤ 0.001). **D** Comparison of *MAPK15* expression levels in patients stratified by fibrosis stage, using classifications from Pantano and coll. (Pantano *et al*., 2021) (ANOVA p = 0.00024). **E** Same as in **D** but using classification of MASLD stages scheme adopted in the work of Govaere and coll. (Govaere *et al*., 2020) (ANOVA p = 0.03504). In each cohort, the ANOVA model compared mean *MAPK15* expression level between disease stage adjusting for the effect of sex. Tukey’s HSD pairwise comparisons between control and pathological groups reach significance only in Pantano cohort (*p ≤ 0.05, **p ≤ 0.01, ***p ≤ 0.001), while in the Govaere cohort they did not reach statistical significance due to the small size of the control group which emphasizes the measured variance.

Based on the histological liver features in WD-fed *Mapk15^-/-^*mice suggesting a progressive MAFLD-like phenotype, we next investigated the presence of specific markers of hepatic damage and inflammation, usually found in human MASH. Indeed, histological examination of their livers showed increased number of lymphocytic infiltrates and frequent “ballooning” injury of the hepatocytes (Kleiner *et al*., 2005) as compared to WD-fed *Mapk15^+/+^*mice (**Fig. 8A**). Interestingly, WD-fed *Mapk15^-/-^* livers also exhibited increased local levels of different proinflammatory cytokines when compared to both WD-fed *Mapk15^+/+^* livers and also to livers from SD-fed *Mapk15^+/+^* and *Mapk15^-/-^*animals, namely TGF-α (Ahmed *et al*., 2022), IL-1α (Wieckowska *et al*., 2008; Gadd *et al*., 2014) and MCP1 (Poulsen *et al*., 2022), while TNF-α levels, although increasing, did not reach statistical significance (**Fig. 8B**). Importantly, *Mapk15^-/-^* liver tissue trichrome stain showed evidence of periportal and perisinusoidal fibrosis compared to WD-fed *Mapk15^+/+^*mice (**Fig. 8C**), overall demonstrating a faster progression of *Mapk15^-/-^* mice to a MASH-like phenotype upon a typical steatogenic insult (i.e., WD), ultimately suggesting a key protective role for this gene against MASLD pathogenetic stimuli. Therefore, since transcriptomics studies of liver biopsies from human MASLD cohorts have already helped to identify specific genes whose altered expression is associated with MASLD progression, to support a protective role of MAPK15 against human MASLD, we evaluated the expression level of this gene using two publicly available patients’ cohorts (Govaere *et al*., 2020; Pantano *et al*., 2021). Transcriptomic data and clinical covariates were collected for a total of 344 patients encompassing various stages of MASLD progression and stages of fibrosis. Our findings revealed that *MAPK15* expression was significantly higher in MASLD patients, compared to healthy controls, in both Pantano (two-way ANOVA p = 0.00024) (**Fig. 8D**) and Govaere (two-way ANOVA p = 0.03504) (**Fig. 8E**) cohorts. This difference, further examined in the Pantano cohort due to a comparable number of patients across groups, remained significant in pairwise comparisons between controls and each stage of fibrosis (**Fig. 8D**).

## DISCUSSION

MASLD is a pandemic metabolic disorder affecting nearly a fourth of worldwide population, with very high human and socioeconomic costs (Loomba *et al*., 2021). A better understanding of the pathogenesis of this disease is therefore extremely urgent, to develop new therapies able to control its long-term deleterious effects. Indeed, while hepatic steatosis is often self-limited in the context of MASLD, under the pressure of environmental factors and/or genetic predisposition, up to 15% of affected individuals can progress and exhibit some degree of MASH, the inflammatory subtype of MASLD (Cohen *et al*., 2011). Alarmingly, up to ∼30% of MASH patients can, in turn, progress to cirrhosis, a serious premalignant condition often (up to 25% of cases) evolving in HCC or liver failure, therefore requiring liver transplantation (Cohen *et al*., 2011). Still, the heterogeneous, fluctuating, and slow natural history of the disease usually requires long-term studies to demonstrate the effects of an intervention on clinical outcomes, and such studies are currently limited. Therefore, preclinical mouse models for MASLD have become crucial tools to investigate mechanisms as well as novel treatment modalities during the progression from steatosis to MASH and subsequent HCC, in preparation for human clinical trials. Still, although there are numerous genetically-, toxins-and diet-induced models of MASH, not all of them faithfully phenocopy and mirror the human pathology very well, leading to huge efforts for defining their metabolic relevance to the human disease (Vacca *et al*., 2024). Among such models, western-type diets, composed of saturated fats, carbohydrates, and 0.1% to 2% cholesterol, akin to fast-food diets, are increasingly recognized as the best models for mimicking human MASH pathology and comorbidities (Vacca *et al*., 2024). Still, while these diets can induce, in mice, obesity with dyslipidemia and elevated plasma liver enzymes and increased liver steatosis with extensive inflammatory infiltration and hepatocyte ballooning (Charlton *et al*., 2011), they often require extended durations to obtain significant fibrosis, especially at low cholesterol concentrations (Vacca *et al*., 2024). Intriguingly, we have shown that, when associated to WD, deletion of the *Mapk15* gene strongly accelerated the insurgence of liver inflammation and fibrosis, closely mimicking the progression of the disease to MASH. This may enhance our ability to study the disease and test behavioural or pharmacological interventions aimed at slowing, halting or even reversing the necro-inflammatory responses triggered by the lipotoxic insult caused by intracellular lipid accumulation (Cohen *et al*., 2011). Furthermore, our mouse model also allows for direct testing of a role for MAPK15 in the progression of MASLD to HCC, the third leading cause of cancer-related deaths and the sixth most common cancer globally (He *et al*., 2023). This could prove to be particularly important because, besides lipotoxicity, HCC is associated with additional important aetiologies traceable to other infective (HBV, HCV) or chemical (alcohol, drugs) causes of inflammatory hepatocyte stress, which could altogether take advantage of specific pharmacological approaches targeting MAPK15 for prevention or cure.

Hepatic steatosis usually arises when hepatic DNL and fatty acid uptake from the circulation saturate the capacity of the liver to oxidize FAs and secrete TG in the form of VLDL. While we have here identified a specific mechanism dependent on MAPK15 depletion causing the increase in CD36 expression and membrane localization, we also excluded a potential role for this MAP kinase in the control of DNL. Importantly, this conclusion is also supported by the fact that DNL is differently regulated in liver and adipose tissue (e.g., adipose lipogenesis is significantly downregulated upon chronic overnutrition and obesity while hepatic lipogenesis is upregulated under similar conditions) (Jeon *et al*., 2023) while we observed increased accumulation of LDs in both tissues, upon *MAPK15* gene deletion, again suggesting that this phenotype is not dependent on DNL. Consequently, we turned our attention to a mechanism similarly regulated in both liver and adipose tissue as cells from both tissues strongly rely on CD36 to uptake FAs and a key role for this protein has been also demonstrated in the pathogenesis of MASLD (Glatz *et al*., 2010; Rada *et al*., 2020).

Fibrosis is the main histologic feature of MASH that predicts clinical outcomes of the disease (Loomba *et al*., 2021). Therefore, accumulation of collagen fibers in WD-fed *Mapk15^-/-^*mice strongly suggests an important role for this MAP kinase in protecting liver from evolution into MASH and, possibly, liver cirrhosis. Since hepatocytes are a rich source of soluble signals driving stellate cell activation, the extremely high levels of accumulated lipids observed in *Mapk15^-/-^* mice suggest that lipotoxicity deriving from this process could be a key factor driving cell injury and consequent release of paracrine signals (ROS, oxidized phospholipids, leptin and different chemokines) with profibrotic activity (Loomba *et al*., 2021). Still, current knowledge on MAPK15 cellular functions also points to different mechanisms working in conjunction with intracellular lipid accumulation to produce the profibrotic effects resulting from the deletion of this kinase. Among such mechanisms, we have in fact recently demonstrated a key role of MAPK15 in the control of intracellular oxidative stress by promoting mitochondrial fitness (Franci *et al*., 2022) and by controlling the activity of the main antioxidant transcription factor, NF-E2-Related Factor 2 (NRF2) (Franci *et al*., 2024). Consequently, its depletion may possibly further worsen oxidative stress in both hepatocytes and stellate cells, exacerbating fibrosis and potentially causing oxidative DNA damage. Moreover, MAPK15 depletion may also induce cellular senescence triggering a senescence-associated secretory phenotype (SASP) (Franci *et al*., 2022), whose chemokine components often have profibrotic activities and may even alter the immune microenvironment and support MASH development or even tumorigenesis (Loomba *et al*., 2021; Meijnikman *et al*., 2021). Ultimately, it was intriguing to observe more subtle differences in the tendency of female *Mapk15^-/-^*mice to accumulate hepatic fat than their wild-type counterparts, although our results were in line with those obtained in most laboratory mouse strains used to model human MASLD by high-fat diets (Norheim *et al*., 2017). In turn, such results suggest that our *Mapk15* knockout model may represent a useful tool also to try to understand underlying hormonal causes for these differences between sexes (Smiriglia *et al*., 2023), which we plan to tackle by next studying hepatic steatosis comparing pre-and post-menopausal and ovariectomized SD-and WD-fed female *Mapk15^-/-^*mice.

Overall, our current work characterizes a new mouse model for progressive MASLD which closely mirrors the human disease, in terms of both histologic and biochemical characteristics, as well as sex-related differences in the severity of their manifestations. Importantly, we also identified a specific mechanism driving this phenotype, based on the CD36-dependent increase in intracellular lipid accumulation, disclosing a potentially novel and important role for a currently undervalued MAP kinase in protecting the liver from an “epidemic” disease for which very few pharmacological approaches are available. Accordingly, our analysis of *MAPK15* expression in livers from human cohorts showed an overall increase in the different stages of MASLD compared to the unaffected individuals, supporting a model in which liver cells induce the expression of this kinase to counteract deleterious effects of lipotoxic stress due to different steatogenic stimuli. Importantly, as preclinical studies have demonstrated that reverting hepatic steatosis is also able to resolve liver inflammation, liver fibrosis, and diabetes, our data provide a new actionable therapeutic target with potential for preventing or reverting MASH and, possibly, contribute to reduce its frequent debilitating and deadly consequences, such as cirrhosis and HCC.

## METHODS

### Mouse models

*Mapk15^-/-^* mice were obtained from Lexicon Pharmaceuticals. In this model, the full coding sequence of *Mapk15* has been substituted with a *LacZ/Neo* gene. The mutation has been generated in 129SvEvBrd strain-derived embryonic stem cells. Chimeric mice have been bred to C57BL/6J albino mice to generate F1 heterozygous animals on 129SvEvBrd/C57BL/6J background. Heterozygous and homozygous mice are vital and with no signs of suffering. Mice used in this study are on a pure C57BL/6J background after crossing into it for at least 10 generations. Inbred C57BL/6J control *Mapk15* wild-type (WT, *Mapk15^+/+^*) mice (JAX stock #000664) were obtained from Jackson Laboratory (Bar Harbor, ME, USA) via Charles River (Calco, LC, Italy) at 7 weeks of age. All mice were socially housed in groups of up to 4 mice per cage under standard conditions (12 h light cycle, ambient temperature of 20-23°C), and free access to food and water. Mice in the WD group were switched from the standard diet [SD: 3% fat, 4RF21, Mucedola, Italy; Kcal 18.5% from protein; 3% from fat; 53.5% carbohydrates (3% sucrose); kcal/g 3.150] to the lipid-rich diet [WD: 0.2% cholesterol and 21% butter, Western U8958 version 35, SAFE®, France; Kcal 14,4% from protein; 38.1% from fat; 47% carbohydrates (33% sucrose); kcal/g 4.2594] starting from 8 up to 24 weeks of age. The WD was stored at 4°C and replaced once per week to avoid lipid peroxidation. All WD-fed mice appeared healthy and active throughout the diet intervention period, and no mouse had to be euthanized before 17 weeks of feeding. Mice in the control group were fed with SD from weaning until 24 weeks of age. Body weight (BW) and food consumption were monitored twice per week, immediately after food replenishment (Ali and Kravitz, 2018). Food pellets were weighed twice per week and the amount of food left in the cages was subtracted from the initially recorded amount. For each mouse, average daily food weight intake per week was calculated. According to NIH-MMPCs guidelines, we calculated the BW change from the initial measurements to analyze the overall effect of diet on this parameter (Ellacott *et al*., 2010). Furthermore, the Body Condition Score (BCS) was monitored at relevant time points (every 8 weeks) over a 17-week period, as recommended (Ullman-Culleré and Foltz, 1999). The food intake of mice was analyzed as the daily amount of food (g) consumed by single mouse for each experimental week. Mice were sacrificed and analyzed at 24 weeks of age. Detailed methodologies for *in vivo* animal procedures have been previously described (Gargiulo *et al*., 2024). Animal experiments were performed in accordance with Directive 2010/63/EU and D.L. 26/2014 and with NIH guidelines for the use and care of live animals and approved by the animal study protocol approved by the Animal Welfare Board of Fondazione Toscana Life Sciences, Siena, Italy, (protocol code: 9AECF.34) and by the Italian Ministry of Health (protocol code: 175/2021-PR).

### Cell Culture

HepG2 Cells were purchased from ATCC, HepaRG were purchased from (Thermo Fischer Scientific, Waltham, MA, USA, Gibco, #HB-8065) (see also **Reagents and Tools section 1.1**). Cell lines were cultured in the adequate medium at 37 °C in an atmosphere of 5% CO_2_/air. Specifically, HepG2 cells were maintained in Earle’s Salts Modified Eagle’s Medium (EMEM -Euroclone S.p.A., Milan, Italy, #ECB3054D), supplemented with 10% heat-inactivated fetal bovine serum (FBS Euroclone S.p.A., Milan, Italy, #ECS0180L), 2 mM L-glutamine (Euroclone S.p.A., Milan, Italy, #ECB3000D), 1x Non-Essential Amino Acids (Merck KGaA, Darmstadt, Germany, Sigma-Aldrich #M7145-100ml), 1 mM sodium pyruvate (Euroclone S.p.A., Milan, Italy, #ECM0542D). HepaRG cells were maintained in William’s E Medium, no phenol red (Thermo Fischer Scientific, Waltham, MA, USA, Gibco, #A1217601) supplemented with 10% heat-inactivated fetal bovine serum (FBS -Euroclone S.p.A., Milan, Italy, #ECS0180L), 2 mM L-glutamine (Euroclone S.p.A., Milan, Italy, #ECB3000D), 5 ug/ml inulin (Merck KGaA, Darmstadt, Germany #I6634) and 50 µM hydrocortisone hemisuccinate (DBA Italia s.r.l, Milan, Italy, MedChemExpress, #HY-B1402).

### RNA interference

MAPK15-specific siRNA (siMAPK15; target sequence 5’-TTGCTTGGAGGCTACTCCCAA-3’) and control non-silencing scrambled siRNA (siSCR, target sequence 5’-AATTCTCCGAACGTGTCACGT-3’) were obtained from Qiagen. MAPK15-specific siRNA #2 (siMAPK15 #2; target sequence 5’-GACAGAUGCCCAGAGAACATT -3’) was obtained from Eurofins Genomics. CD36-specific siRNA was purchased from Santa Cruz Biotechnology (#sc-29995). Lipofectamine RNAiMAX (Thermo Fischer Scientific, Waltham, MA, USA, Invitrogen, #13778075) reagent was used specifically for the delivery of siRNA with reverse transfection as manufactory protocol. Briefly, for transfection of six wells plate, for each well, 60nM of siRNAs, 7,5 µl of RNAiMAX and 500 µl of Opti-MEM (Thermo Fischer Scientific, Waltham, MA, USA, Gibco, #11058021) were added to wells and incubated for 20-10 minute respectively for HepG2 and HepaRG. Then, 1,4 x 10^5^ of HepG2 cells or 1,7 x 10^5^ of HepaRG, were seeded on the complexes. Samples were collected 72 h after transfection. When indicated, co-transfection of siRNAs was performed.

### Antibodies

Primary and secondary antibodies are listed in **Reagents and Tools section 1.2**

### Western blots

Cells were washed with cold PBS and total protein extracts obtained by adding “RIPA buffer” (50 mM Tris-HCl pH=8.0, 150 mM NaCl, 0.5% Sodium deoxycholate, 0.1% SDS, 1% NP-40, 2 mM Orthovanadate, 2mM Sodium fluoride, 1mM Dithiothreitol and 1× protease inhibitors) or “1% SDS hot lysate buffer” (10mM Tris-HCl pH=8.0, 1% SDS, mM Na-Orthovanadate, ddH_2_0). Following mechanical detachment using cell scrapers, total lysates were gathered in tubes, subjected to vortexing, and then incubated for 15 min on ice, while for the 1% SDS hot lysates were incubated at 90°C for 15 min (vortexing every 5 min), followed by ultrasonic cell disruptor (on ice) to break all cell clusters until the lysate becomes clear. Livers from mice were collected and total lysates were obtained as described before (Gherardini *et al*., 2021). Briefly, ∼30mg of each sample were disrupted by bead homogenization in Lysing Matrix D tubes using a FastPrep-24 5G homogenizer (MP Biomedicals, Solon, OH, USA) filled with 600 µl of RIPA or 1% SDS hot lysate buffer (1:20 w:v). Next, samples were transferred in new tubes and then incubated as previously described. All protein samples were centrifuged for 10 min at 16,000 g, and supernatants were collected, representing total cell lysates.

Total proteins were quantified by Bradford assay and the same amount of protein extracts was used for western blot analysis. Laemmli Loading Buffer 5X (250 mM, Tris-HCl pH 6.8, 10% SDS, 50% glycerol, bromophenol blue) was added to protein samples at 1X concentration, which were then heated for 5 min at 95°C. Samples were loaded in 8% acrylamides gel and resolved by SDS-PAGE, transferred to Immobilon-P PVDF membrane (Merck KGaA, Darmstadt, Germany, Millipore, #IPVH00010). Membrane was blocked 2 h with 5% LFDM (GeneSpin Srl, STS-M500) in TBST at RT probed with the indicated primary antibody overnight (ON, 16 h), washed 3 times with 5% LFDM and probed 1 h with the appropriate secondary antibody, than washed 2 times with 5% LFDM, 3 times with TBST, and revealed by enhanced chemiluminescence detection (ECL Plus; GE Healthcare, Chicago, IL, USA, #RPN2132). Densitometric analysis of western blots was performed with NIH ImageJ (National Institutes of Health).

### Immunofluorescence and confocal microscopy

Seven × 10^4^ HepG2 and 6 × 10^4^ HepaRG cells were seeded in 12-well cell culture plates and transfected with each siRNA as described. For confocal analysis, cells were washed with PBS, fixed 4% paraformaldehyde (PFA) for 20 min and then blocked for 30 min with 1% BSA in PBS. Cells were then incubated with appropriate primary antibodies for 30 min, washed three times with PBS and then incubated with secondary antibodies for 30 min with 2% BSA in PBS. Where indicated, LDs were stained using 3.6 μM of BODIPY 493/503 (Cayman Chemical, #25892). Nuclei were stained with 6 μM 4′,6-diamidino-2-phenylindole (DAPI) in PBS for 10 min. Coverslips were mounted in a fluorescence mounting medium (Dako, S3023) and stored at 4°C. After 24 h samples were visualized on a TSC SP5 confocal microscope (Leica, #5100000750) installed on an inverted LEICA DMI 6000CS (10741320) microscope using an oil immersion PlanApo 40× 1.25 NA. Images were acquired using the LAS AF acquisition software (Leica) and LDs count or intensitometric analysis of fluorescence were performed through unbiased analysis using the Quantitation Module of Volocity software (PerkinElmer Life Science, I40250). Data were obtained by analyzing at least 400 cells from three different experiments.

### Cytofluorimetry

Eight × 10^4^ HepG2 and 1 × 10^5^ HepaRG cells were seeded in 12-well cell culture plates and transfected with each siRNA as described. For cytofluorimetric analysis cell were detached and washed with PBS, then fixed 1,5% PFA for 10 min and blocked for 30 min with 2% BSA in PBS. Next, samples were fixed again with 1.5% PFA for 5 min, washed and then incubated with secondary antibodies for 30 min with 2% BSA in PBS, then were washed, resuspended in PBS and acquired. For LDs analysis, cells were detached and washed with PBS, then fixed 1.5% PFA for 10 min. LDs were stained using 3.6 μM of BODIPY 493/503 (Cayman Chemical, #25892) for 30 min, then were washed, resuspended in PBS and acquired. Samples were acquired on NovoCyte Quanteon Flow Cytometer Systems, data were analyzed with FlowJo software. All analyses were performed in triplicate.

### Fatty Acid Uptake

Palmitate cellular uptake was evaluated by measuring the relative fluorescence intensity in cells after incubation with BODIPY-palmitate (BODIPY-PA, Cayman, #26749). Briefly, HepG2 and HepaRG cells were cultured and transfected with appropriate siRNA. After 72 h, cells were incubated with complete medium without FBS and 200 µM Sulfo-N-succinimidyl oleate (SSO, MedChemExpress, #HY-112847A) (Jay *et al*., 2020) or DMSO as a control, for 6 h. Next, cells were incubated in 2.0 ìM BODIPY-PA at 37°C in an atmosphere of 5% CO_2_/air for 30 min. After washing, the samples were detached, fixed by 1,5% PFA for 10 min, washed and analyzed on a NovoCyte Quanteon Flow Cytometer Systems. Data were analyzed with FlowJo software. All experiments were performed in triplicate.

### Analysis of gene expression

Total Cell lines RNA lysates were obtained using QIAzol Lysis Reagent (Qiagen, Hilden, Germany #79306) followed by RNA purification. Livers from mice were collected and ∼30mg of each sample were disrupted by bead homogenization in Lysing Matrix D tubes using a FastPrep-24 5G homogenizer (MP Biomedicals, Solon, OH, USA) filled with 1ml of QIAzol Lysis Reagent (Qiagen, Hilden, Germany #79306) before RNA purification. Reverse transcription was performed with the QuantiTect Reverse Transcription Kit (Qiagen, Hilden, Germany #205311). RT-qPCR was performed with Luna Universal qPCR Master Mix on a Rotor-Gene 6000 RT-PCR system (Corbett Life Science). Primer pairs were purchased from Eurofins Genomics (Ebersberg, Germany). Their sequences are reported in **Reagents and Tools section 1.3.**

### Luciferase Assays

HepG2 cells were seeded at 6 × 10^4^ cells/well density in 12-well plates, in triplicate, and were reverse-transfected with siSCR and siMAPK15. After 48-h of RNA interference, cells were transfected with 500 ng of pSynSRE-T-Luc was a gift from Timothy Osborne (Addgene plasmid # 60444; http://n2t.net/addgene:60444; RRID:Addgene_60444).After 24 hours, samples were lysed in Passive Lysis Buffer (Promega, Madison, WI, USA) and processed as indicated by the manufacturer. Luciferase activity in cellular lysates was assessed on a Glomax 20/20 luminometer (Promega, Madison, WI, USA) using the Luciferase Assay System (Promega, Madison, WI, USA). Results were normalized for cell count. All samples were read at least in triplicate.

### Statistical Analysis

Significance (p-value) was assessed, where indicated, by either Unpaired t test with Welch’s correction, Two-way ANOVA-Mixed effect model - Repeated measures, One-way ANOVA – Mann Whitney nonparametric test, using GraphPad Prism8 software. Asterisks were attributed as follows: *p ≤ 0.05, **p ≤ 0.01, ***p ≤ 0.001, ****p ≤ 0.0001.

### Histology and immunohistochemistry sample preparation

Immediately after removal, mice samples were formalin-fixed and paraffin-or OCT-embedded as previously described (Gargiulo *et al*., 2024). Briefly, 7 ìm serial sections were cut and stained with haematoxylin and eosin (H&E) for morphological evaluation. Masson’s trichrome and Oil-Red O staining were used to assess collagen and lipid content, respectively. Paraffin-embedded liver sections were used for Masson’s trichrome staining following manufacturer’s instructions (Abcam, Cambridge, United Kingdom). OCT-embedded liver sections were formalin-fixed for 2 min, and Oil-Red O stained to identify LDs in hepatocytes. Each slide was incubated for 15 min in 0.3% Oil Red O in isopropanol and counterstained with Carazzi’s haematoxylin.

Paraffin-embedded liver sections were used for immunohistochemistry with CD36 antibody. Heat-induced epitope retrieval (HIER) was performed at 98°C for 10 min in 10 mM citrate buffer, pH6. Slides were then incubated with the CD36 antibody ON at 4°C (dilution 1:200), and then with the appropriate alkaline phosphatase conjugated secondary antibody following manufacturer’s instructions (Vector Red Substrate kit, alkaline phosphatase, #SK-5100). All sections were counterstained with haematoxylin. The slides were observed with the Nikon Eclipse E600 optical microscope. Images were acquired with the Nikon DS-U1 camera and morphometric analysis was performed with the Nis Elements v3AR software (Nikon Instruments). Specifically, liver steatosis and inflammation were assessed on H&E-stained slides. At least five microscopic fields per section (400× magnification) and three sections per animal were examined. The grade of fibrosis was assessed on Masson’s trichrome-stained sections by measuring the area covered by blue staining as a fraction of the total analyzed area. Data are reported as fold changes. H&E visceral adipose sections were used to measure the cross-sectional area of at least 20 adipocytes per animal.

### Radioactive assay

HepG2 cells were seeded and interfered for *MAPK15* expression into 12-well plates. The incorporation of ^14^C-glucose into lipids was performed as previously described (Lorito *et al*., 2024). Briefly, after 56 hours of MAPK15 downregulation, culture media were supplemented ON with 1 mCi of the radiolabeled metabolite (Perkin Elmer, Waltham, Massachusetts, United States, #NEC042V250UC). Cells were next washed in ice-cold PBS and lysed in RIPA buffer. Lipids were extracted resuspending in 4 volumes of a CHCl_3_:MeOH (1:1). The solution was then centrifuged at 2000 *g* for 5 min at room temperature. The lower phase containing lipids was collected, transferred to a scintillation vial, and the incorporated radioactive metabolite-derived signal was measured on the scintillation counter and normalized to cell count.

### Biochemical analysis

At 24 weeks of age, mice were fasted for 3 hours, and blood samples were collected by cardiac puncture (40-60 seconds) prior to euthanasia under 3% isoflurane + 2 lt/min oxygen anaesthesia. Blood samples were centrifuged at 3500 rcf for 15 min to obtain sera, which were immediately frozen (−20°C). Serum alanine aminotransferase (ALT), aspartate transaminase (AST), cholesterol, triglycerides, glucose, results were provided by experienced laboratories (IZSLT, Rome, Italy; Galileo Research, Pisa, Italy) accredited by the Italian Ministry of Health to carry out studies on rodents for research purposes. In addition, fasted serum insulin was determined using an Ultrasensitive mouse insulin ELISA kit (Mercodia, Uppsala, Sweden, 10-1249-01) according to the manufacturer’s instructions. Insulin sensitivity was assessed by using the homeostatic model assessment for insulin resistance (HOMA IR) method (Fraulob *et al*., 2010).

### Detection of local cytokine release

The total liver protein extract was analysed to determine the concentrations of cytokines. TGF-β, IL-1β, MCP1 and TNF-α were measured by a custom MILLIPLEX MAP High Sensitivity Human Cytokine Magnetic Bead Panel (Merck KGaA, Darmstadt, Germany, Millipore) based on the Luminex xMAP Technology (Luminex Corporation, Austin, Texas, United States) as described on manufactory protocols. Data were read on the Luminex MAGPIX Instrument (Luminex Corporation, Austin, Texas, United States) and analysed using the MILLIPLEX Analyst 5.1 software (Merck KGaA, Darmstadt, Germany, Millipore). Concentrations of cytokines were calculated using a standard curve and normalized to the protein concentration.

### Analysis of MAPK15 expression in human patient datasets

To investigate the expression of *MAPK15* in the various stages of the MASLD, we collected two cohorts from Pantano et al. (N=143, GSE162694) (Pantano *et al*., 2021) and Govaere et al. (N=201, GSE135251) (Govaere *et al*., 2020). Specifically, we collected and analysed the release of expression counts aligned against the GRCh38.14 by the NCBI

To minimize potential biases introduced by differences in RNA-seq experimental procedures and protocols, each cohort was analyzed separately. Raw counts were normalized using the Trimmed Mean of M-values (TMM) method, followed by the application of the voom transformation from the limma package (Ritchie *et al*., 2015). To assess differential expression of *MAPK15* between groups, we employed an ANOVA model that corrects for the effect of sex. Post-hoc analyses were conducted using Tukey’s Honestly Significant Difference (HSD) test to explore within-group differences under the same model, ensuring robust pairwise comparisons. All the analysis was conducted using R (v. 4.4.1).

## Abbreviations

ACC1: Acetyl-CoA Carboxylase 1
ALT: Alanine aminotransferase
AST: aspartate aminotransferase
BW: body weight
BCS: Body Condition Score
BW: body weight
CD36: Cluster of Differentiation 36
ChREBP: Carbohydrate-responsive element-binding protein
DNL: *de-novo* lipogenesis
ER: endoplasmic reticulum
FAs: fatty acids
FAO: FAs β-oxidation
FASN: Fatty Acid Synthase
FER: food efficiency ratio
HCC: hepatocellular carcinoma
HFD: high-fat diet
IHC: immunohistochemistry
HOMA IR: homeostatic model assessment for insulin resistance
HRI: hepatorenal index
LDs: lipid droplets
LW: liver weight
KO: knockout
LCFA: long chain fatty acids
MAFLD: metabolic dysfunction-associated fatty liver disease
MASL: metabolic dysfunction-associated steatotic liver
MASH: metabolic dysfunction-associated steatohepatitis
MASL: metabolic dysfunction-associated steatotic liver
MASLD: metabolic dysfunction-associated steatotic liver disease
MAPK15: mitogen activated protein kinase 15
NAFLD: non-alcoholic fatty liver disease
OA: oleic acid
PA: palmitic acid
RT-qPCR: quantitative reverse transcription polymerase chain reaction
SCD1: Stearoyl-CoA Desaturase-1
SRE1: sterol regulatory element-1
SREBP: sterol regulatory element binding protein
SD: standard diet
SSO: Sulfo-N-succinimidyl oleate
TAG: triacylglycerol
TG: triglyceride
T2D: type 2 diabetes mellitus
VLCFA: very-long chain fatty acids
VLDL: very-low-density lipoproteins
WD: Western diet
WT: *wild-type*

## ACKNOWLEDGEMENTS

We thank Federico Galvagni (University of Siena) and Francesca Carlomagno (University of Napoli) for their critical reading of the manuscript. This research was supported by Next Generation EU [DM 1557 11.10.2022] to MC and MGA, in the context of the National Recovery and Resilience Plan, Investment PE8 – Project Age-It: “Ageing Well in an Ageing Society”. The views and opinions expressed are only those of the authors and do not necessarily reflect those of the European Union or the European Commission. Neither the European Union nor the European Commission can be held responsible for them. This work was also supported by Ministero dell’Istruzione dell’Università e della Ricerca – Progetto di Ricerca di Rilevante Interesse Nazionale (PRIN) 2022RCFZZ3 to AM.

## DATA AVAILABILITY STATEMENT

The data associated with this paper and further information and requests for resources and reagents should be directed to and will be fulfilled by the lead contact, MC.

## AUTHORS’ CONTRIBUTIONS

Conceived and designed the project: MC. Supervised the project: EB, RDA, MGA, GB, AG, AM, VB, MC. Performed most of the experiments and analyzed data: GI, SG, DB, LG, NL, SDT, DT. Helped in experiments and data analyses: LF, TT. Wrote the manuscript: MC. Edited the manuscript: GI, SG, LG, LF, NL, RDA, MGA, GB, AM, VB, MC. Discussed and interpreted the results and approved the manuscript: all authors.

